# Anesthesia-related brain microstructure modulations detected by diffusion MRI

**DOI:** 10.1101/2022.10.04.510615

**Authors:** Thomas Beck Lindhardt, Leif Østergaard, Zhifeng Liang, Brian Hansen

## Abstract

Brain clearance has been found to be greatly enhanced during sleep and anesthesia. Studies using optical microscopy have attributed this to an anesthesia-related expansion of the brain’s extra-cellular space. These results, however, have been based on invasive experiments with a limited field of view. Here, we employ diffusion kurtosis magnetic resonance imaging to assess brain microstructure in the awake and anesthetized mouse brain. We find both mean diffusivity and mean kurtosis to be significantly decreased in the anesthetized mouse brain compared to the awake state. This effect is observed in both gray matter and white matter. Our findings are consistent with the reports of brain ECS volume increase in sleep and anesthesia. Based on simple simulations, we discuss how an ECS volume increase (cell shrinkage) during sleep and anesthesia can coexist with other aspects of brain physiology. Our study demonstrates that diffusion kurtosis MRI can be used in the study of the glymphatic system. Importantly, our study shows an overlooked effect of anesthesia on brain microstructure which is relevant to preclinical MRI as a whole.

## 1 Introduction

Removal of toxic waste solutes from the brain’s extracellular environment is essential for brain homeostasis and health. It is still unclear how exactly these waste products are cleared from the brain; however, compelling evidence is now accumulating and points towards a novel, true circulatory ‘glymphatic’ system capable of providing nutrients and clearing waste solutes simultaneously. In its original form as proposed in 2012^1^, the glymphatic system is a brain-wide network of perivascular channels, formed by glial cells surrounding the arteries and veins, that enable cerebrospinal fluid (CSF) to travel along penetrating arteries, where it would, by flow mechanisms, enter the extracellular space (ECS) allowing for interchange with the interstitial fluid (ISF)^2^. The exchange between para-arterial influx of CSF and para-venous efflux of ISF would then allow brain fluid and waste solutes, such as Alzheimer-related Aβ42-proteins to exit the extracellular space (ECS), before later being transported out of the brain to larger lymph vessels. For this reason, the glymphatic system and its function has been suggested to play a crucial role in safeguarding the brain from neurodegenerative diseases such as Alzheimer’s disease (AD). Intriguingly, a follow up study found that the rate of glymphatic clearance was greatly enhanced during sleep and under anesthesia as compared to wakefulness^3^. In their study using 2-photon microscopy (2PM), cisterna magna infused CSF tracers spread faster and covered a 60% larger area of the visible parenchyma in sleeping and anesthetized mice compared to awake mice. This was attributed to a corresponding increase in ECS volume and validated by real-time tetramethylammonium iontophoresis (TMA^+^) quantification. The explanation offered was that such ECS expansion would presumably decrease the total tissue resistance of fluid flow in the ECS allowing for a more rapid para-arterial influx of CSF and facilitate greater fluid movement in the brain parenchyma.

Indeed, the original glymphatic hypothesis is compelling due to its simplicity and it provides potential answers to long standing questions on how solutes and fluid pass through the brain parenchyma. It has, however, been intensely debated, especially due to that fact that much of the work preceding the generation of the glymphatic hypothesis, is based upon data from different microscopy modalities with access to only a small part of cortex and from that generalized to whole brain functionality. It has now been found that flow mechanisms alone cannot drive fluid transport within the interstitial spaces of brain parenchyma and it is now accepted that diffusion is the dominant process accounting for solute transport in the brain parenchyma although flow components may still exist^4^. For example, computational models show that the flow rate required to surpass that of diffusion in parenchyma created by a hydrostatic pressure would be 40 times that of the CSF production rate of the choroid plexus (CP)^5,6^. A comprehensive description of the entire theory, controversies and current evidence regarding all aspects of the glymphatic hypothesis is beyond the scope of this article, however please see excellent reviews^4,7,8^. Alongside these and other aspects concerning the extravascular solute transport system, as proposed in the original glymphatic hypothesis, the phenomenon of a 60% increase of ECS volume during sleep or anesthesia has also been discussed extensively^4,7,8^. In normal brain tissue the ECS volume fraction (α) is typically estimated at 20% under normal physiological conditions although α evidently varies throughout different regions of CNS due to the difference in tissue compositions, complexity and cell types. So far, vast changes in α have predominantly only been associated with maturation, aging and different pathological conditions^9^. For example, under global ischemia α is reduced to only 5% while it has been shown that α is reduced by 10-20% in a transgenic mouse model of AD^10^ and during epileptic seizures, α is reduced by 35%^11^. Few studies have investigated the effect of arousal and anesthesia on ECS volume dynamics and so far, results are inconsistent. As mentioned, Xie et al. used TMA^+^ to find that the ECS volume was increased by ∼60% in mouse gray matter both under natural sleep and anesthesia. Similarly, a mouse study using TMA^+^ and sleep-/wake-inducing artificial CSF (aCSF) mixtures, also detected an increased ECS volume in the awake-state compared to sleep, however the effect was much smaller (∼20%)^12^. Conversely, a study using *ex vivo* brain slices of rat visual cortex found that the β-Adrenergic receptor agonist DL-isoproterenol (ISO) deceased α^13^. Another study in rats also found that gas isoflurane induced a decrease in the ECS volume when compared to intravenous injections of dexmedetomidine and sodium pentobarbital anesthesia^14^. In addition to the different TMA^+^ studies investigating the dynamic changes of ECS parameters during arousal and anesthesia, several groups also applied magnetic resonance imaging (MRI) as a means to cover the whole brain while keeping the skull intact. For example, Gakuba et al.^15^ used diffusion-weighted magnetic resonance imaging (dMRI) to obtain global apparent diffusion coefficient (ADC) maps. The ADC reports on overall tissue diffusivity across all compartments (along a single encoding direction) and is known to be sensitive to brain microstructure changes affecting the ECS^16^. The ADC would therefore be expected to be sensitive to the ECS volume changes observed in sleep/anesthesia. However, in their study no significant difference in global brain ADC was found between awake and anesthetized mice. Similarly, Demiral et al.^17^ compared ADC maps obtained from the awake and sleep state using human subjects and did also not detect any global changes in ADC values. Therefore, it remains to be seen if the ECS modulation reported in Xie et al. is detectable with non-invasive methods suitable for whole brain assessment. This is important not only because the existence of sleep-related ECS volume modulations needs to be verified with a range of methods including non-optical methods. Furthermore, non-invasive methods capable of detecting ECS modulations are needed to investigate the spatial and temporal dynamics of such ECS modulations in the living human brain because their function is poorly understood and the physiological details of such a mechanism are unclear. Of particular importance is whether the ECS volume fraction modulations occur in an equal or differentiated manner across different types of brain tissue and cellular compartments.

Lack of sensitivity may be the reason for the negative findings in Gakuba et al.^15^ and Demiral et al.^17^. While the ADC obtained from dMRI is a valid method for estimating free water diffusivity in the whole brain it might not be sufficiently sensitive to detect anesthesia/sleep-related ECS volume modulations. Instead, the dMRI method called diffusion kurtosis imaging (DKI) might be used^18^. Compared with diffusion tensor imaging (DTI), DKI is a refined method for obtaining diffusion-based metrics with increased sensitivity to microstructural changes^19^. The improved sensitivity that DKI offers over DTI, is primarily due to the fact that it accounts for the non-Gaussian diffusion behavior that is found in the complex brain tissue^20^. Consequently, DKI provides an indirect probe of tissue microstructural composition and compartmentalization. DKI yields many parameters that can be mapped on a voxel-by-voxel basis. Several of these DKI metrics have been shown to detect subtle changes in brain tissue structure meaning that DKI is of value both as a diagnostic tool but also for basic neuroscience, e.g. in the study of rapid brain plasticity^21^. Although DKI is very sensitive to microstructural changes, metrics derived from DKI are unspecific. Therefore, when interpreting DKI findings, additional validation is often required by means of either histological comparisons or modelling^22,23^. The ECS volume changes reported in Xie et al. would naturally affect both intracellular space (ICS) volume fraction and other features of the tissue microstructure such as restriction lengths. These effects to both tissue compartmentalization and cell size distribution are likely to be detected by DKI as the method is sensitive to exactly these features of the tissue microstructure.

The primary aim of this study was therefore to investigate if shifts in brain tissue microstructure between the awake and anesthetized state can be detected with DKI. We hypothesized that if the extracellular space volume does indeed change from awake to anesthetized state, the mean diffusivity (MD) and mean kurtosis (MK) parameters obtained from DKI measurements, would be able to detect such changes. Based on previous reports we expect the effect to be present in gray matter (GM), but as MRI offers whole brain coverage, we also analyze white matter (WM) which was not explored in the previous reports using optical imaging techniques. Finally, to aid our interpretation of the obtained DKI results, we use numerical simulations based on MRI microscopy and compartment-based DKI modelling to simulate the effect of ECS volume fraction changes to MD and MK in GM.

In our study, we show that isoflurane anesthesia significantly decreases group mean MD and MK values in GM when compared to the awake state (MD, *P* < 0.05, MK, *P* < 0.001 paired; two-sample Student’s t-test). Similarly, WM displays significant changes to MD and MK (MD, *P* < 0.05, MK, *P* < 0.05; paired two-sample Student’s t-test). Our data supports previous findings that microstructural modulations occur between the awake and anesthetized state. Based on simple DKI compartment modeling, we argue that the observed changes in GM can only occur if cell-volume changes occur in a differentiated manner meaning that different cell types respond with different cell-volume changes which together create the ECS volume modulation observed in previous studies. This would mean that neurons, glia and pericytes may modulate their volume to different extents during anesthesia. Finally, aspects of the potential physiological implications of this are discussed.

## 2 Methods and materials

### 2.1 DKI Theory

Diffusion MRI (dMRI) is sensitive to tissue organization on the cellular level. An increasingly popular strategy to obtain improved microstructural sensitivity in dMRI is the DKI framework^18^ which is an extension of the diffusion tensor imaging (DTI) framework and improves the DTI framework by taking into account the leading deviation from Gaussian diffusion. This deviation is caused by the tissue microstructure influencing the water diffusion profile. DKI is sensitive to microstructure generally^19^ and in brain DKI can be used to investigate both gray matter (GM) and white matter (WM)^24^. The central signal equation in DKI is:

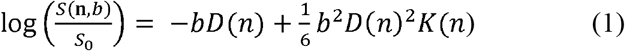

where *b* denotes the b-value, *D*(n) and *K*(n) is the observed diffusivity and kurtosis along encoding direction *n. S*_0_ is the unweighted signal used for normalization. Central DKI parameters are the mean diffusivity (*MD*) and mean kurtosis (*MK*):

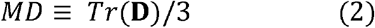

where **D** is the diffusion tensor and Tr is the trace. The MK is defined as:

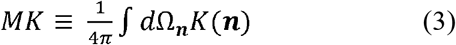

where the integral is over the sphere. These two parameters are also the central parameters investigated here. MD can also be estimated from the simpler DTI framework but studies have shown that DTI parameter estimation is improved within the DKI framework^25^. While both MD and MK have been shown to be sensitive to microstructural remodeling in brain tissue^21,26^ the interpretation of how they reflect specific tissue properties remains a challenge. Of relevance to our study is the relation between MK and tissue compartmentalization as the ECS has been reported to expand in natural sleep and in the anesthetized state^3^. This ECS increase is believed to happen at the expense of ICS volume. It is unclear, however, if all IC compartments (i.e., all cell types) contribute equally to produce this ECS expansion or if some cell types respond differently than others. We will use simple compartment model simulations to shed light on this effect. Direct relations between the apparent kurtosis and compartmental diffusivities are given in^27^:

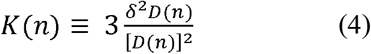

where *δ*_2_*D*(*n*) is the variance of the diffusivity. So, for a system comprised of *N* compartments:

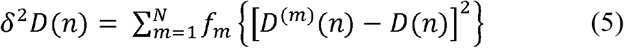

with *D*_(m)_(*n*) being the apparent diffusivity of the *m-*th compartment and the total apparent diffusivity of the system *D*(*n*) is simply the sum of these over all compartments weighted by their volume fractions *f*:

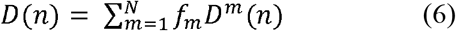

We will use this simple framework to interpret our DKI findings with regard to compartmental size changes between awake and anesthesia. Since the ECS dilation effect reported in Xie et al^3^ is found GM, this tissue is the primary focus of our analysis. This simplifies matters as GM can be considered (near) isotropic meaning that we can assume *MK* = *K*(n) and MD = *D*(n).

### 2.2 Simulations

For our first round of simulation the tissue is just seen as effectively two compartments as is often done in simple dMRI modelling^27,28^. Here, we use the compartmental diffusivity estimates from Maier and Mulkern^28^ to define a two-compartment system with volume fractions *f*_i_ and diffusivities *D*_i_: *f*_1_ = 0.51, *D*_1_ = 0.5 µm^2^/ms; *D*_2_= 1.5 µm^2^/ms. Based on MR microscopy studies of diffusivities in mammalian tissues we assign a faster diffusivity to the ICS than the ECS in keeping with the observation that the effective diffusion coefficient in ECS is only two-fifths that of free diffusion^18^. Using these parameters with Eqs. (4)-(6) yields MD and MK estimates for such a system. In our simulations we assume compartmental diffusivities to be unchanged by any volume fraction modulation meaning that in the simulations *D*_1_ and *D*_2_ are constants. The variation in MD and MK produced by volume fraction shifts can be studied and compared to our data from the awake and anesthetized state. In a second round of simulations, we add another ICS compartment to explore the effect of ECS modulation in a system with different intra-cellular space (ICS) volume shifts among cell types. For this we use values similar to the first round of simulations but with a slight offset between ICS diffusivities: As in our first simulations the first compartment is assigned a diffusivity *D*_1_ = 1.5 µm^2^/ms (ICS_1_) and a volume fraction of *f*_1_ = 1-*f*_2_-*f*_3_ where *f*_2_ is the volume fraction of a compartment with a diffusivity of *D*_2_=0.6 µm^2^/ms (ICS_2_), and *f*_3_ is the size of a compartment with *D*_3_= 0.5 µm^2^/ms (ECS). These simulations therefore differ from the first simulations only in the addition of a second ICS compartment with an intermediate diffusivity. The values used are listed in not meant to capture all aspects of the diffusion properties in tissues and the conclusions drawn do not depend on the exact values used here. This framework allows us to simulate how MK might be expected to change in response to a differentiated change in ICS volumes. By allowing volume fractions *f*_1_ and *f*_2_ to vary independently between 0.1 and 0.45 we can map out the values of MD and MK expected for such a system. This simple model will be used to provide a qualitative interpretation of our DKI results.

### 2.2 Animals

Eight-week-old male and female young adult C57BL/6 mice (Taconic Bioscience Inc., Ejby, Denmark) were used in this study. When mice first arrived at our facility, they were divided sex-wise into groups of two or three and were given two weeks of stable acclimatization. Stable lighting was held on a 12-hour dark/light cycle with lights on from 5:00 am to 5:00 pm and maintained at ambient temperature of 21□±□2 °C while humidity was controlled at 45 ±□5 %. All mice had access to chow and water *ad libitum*. Animal handling and awake MRI habituation was performed by a single experimenter to minimize stress induced by multiple handlers. Throughout the experimental period body weight was monitored to ensure weight loss did not exceed 20% of normal bodyweight. After the last scan animals were deeply sedated with isoflurane (5%) and euthanized with barbiturate (Pentobarbital, 1300mg/kg). All animal housing, handling, and experimental protocols were conducted according to the regulations of the Danish Ministry of Justice and Animal Protection Committees under permit no. 2019-15-0201-00285.

### 2.3 Head holder implantation

The head holder, design and surgical procedure were similar to our previous study^31^. All animal surgeries were performed by one experimenter to minimize variance and conducted under normal aseptic conditions. Briefly, mice were anesthetized with isoflurane (5%) and then mounted on a stereotaxic frame with isoflurane anesthesia maintained with a mixture of isoflurane (1.2-1.5 %), medical air (0.4 mg/l) and oxygen (0.4 mg/l). The complete description of the surgical procedure for head holder implantation can be found in our previous work^31^. In the first four days of the post-surgery period mice received IP injections of antibiotics; 0.2mg/g (STADA, Ampicillin, Vnr: 108259) and inflammatory drugs; 0.1mg/g (ScanVet, Carprofen, Vnr: 027693). Mice also received analgesics 0.12mg/g (Indivior, Temgesic, Vnr: 521634) post-surgery and was administered SC. For the remaining four days analgesics were diluted in the drinking water of their homecage (2 ml Temgesic/120 ml water). After the head holder had been surgically affixed, mice were afforded ten days of recovery, before proceeding with the awake MRI habituation.

### 2.4 Animal handling and awake MRI habituation protocol

To familiarize animals with the experimenter animals were handled for 20 minutes each day on the last five days of the recovery period. During the handling procedure mice were gently picked up and allowed to roam the hands and the lap of the experimenter. Mice were picked up using the tunnel method as it has previously been shown to reduce handling stress^32^. Before transporting the mice back to their homecage they received a drop of condensed milk as reward. After the recovery and handling period, mice were then habituated to the scanner environment for ten consecutive days using a mock MRI setup that as previously described^31^. The mock MRI setup could accommodate three mice simultaneously. Male and female mice were not mixed during the habituation session. On the first three days of habituation mice were head-restrained for 20, 40 and 60 minutes, respectively, and no MRI sound was played. On the fourth day mice were also restrained for 60 minutes but was then introduced to the MRI sound. The following days 20 minutes were added each day up to a total of 120 minutes.

The detailed description of the habituation protocol can be found in (**Fig. 1**). All awake MRI habituation was conducted in the time period from 12:00 am – 14:30 pm in order to minimize influence of circadian rhythms. The estrus cycle of female mice was not taken into account.

**Fig. 1.**
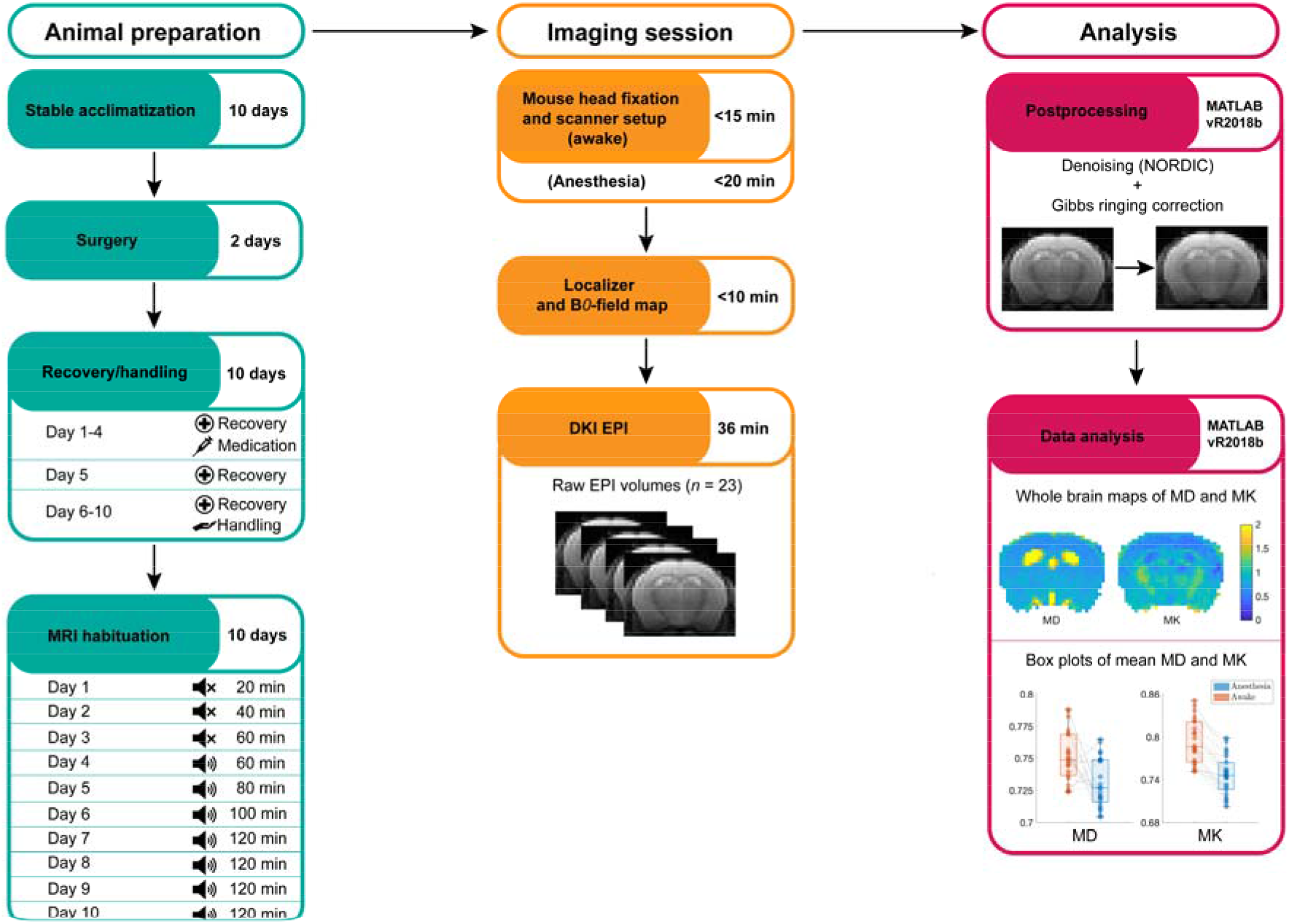
Flowchart of experimental procedures. Description of the different stages of the MRI experiments of head fixed mice in the awake and anesthetized state. Animal preparation (green boxes) describes all the procedural steps and the duration of each step. During the recovery/handling period, mice were given daily dosages of medication (day 1-4). The following day mice were left untouched (day 5), while on the remaining days each animal was handled for 20 minutes per day (6-10). During MRI habituation animals were restrained in the simulation chamber. Twenty minutes were added to the total habituation time each day up to a total duration of 120 minutes. MRI acoustic noise during habituation was added starting from day 4 as indicated by the speaker icons in the bottom box in the first column. Imaging session (orange boxes) lists each step of the MRI setup, the MRI protocols used and the total scan time for each MRI protocol. The time spent on animal setup before scanning commenced was roughly 15 minutes for the awake state and 20 minutes for the anesthetized state. dMRI EPI volumes were acquired for each mouse (*n* = 23) in both awake and anesthetized state. The imaging sessions for awake and anesthesia were conducted one day apart. Analysis (red boxes) describes each of the postprocessing steps and the data analysis. Postprocessing was conducted using NORDIC denoising technique^29^ and Gibbs ringing correction^30^. DKI data processing yielded whole brain maps of MD and MK which were used for further analysis and comparison between groups.

### 2.5 In vivo MRI

MRI data was acquired using a Bruker Biospec 9.4T MRI system equipped a 76 mm volume coil (Bruker Biospin, Germany) for transmission and a high-sensitivity rodent cryo-array surface coil (Bruker Biospin, Germany) for reception. Mice were head fixed with plastic screws onto the 3D printed MRI animal bed and the bed was secured into the bottom of the MRI cradle with another set of plastic screws. The cradle was then gently pushed inside the MRI bore so the head of the animal was positioned directly beneath the surface coil. For anesthesia, mice were briefly sedated with a mixture of isoflurane (5%), medical air (0.4 mg/l) and oxygen (0.4mg/l), while during maintenance isoflurane was decreased to (1.2-1.4%). The custom animal bed for anesthesia was designed with integrated tubing for hot water to ensure stable body temperature which was kept at 37ºC ± 0.5ºC for the duration of the scan. For both scan sessions (awake and anesthesia for each animal), localizer scans were performed to ensure animal positioning and then a B0-field map was acquired (TE 1.84 ms, echo spacing 3.6 ms, TR 20 ms, field of view 30 × 30 mm, matrix size 128 × 128). Last, DKI data was acquired using a diffusion weighted segmented EPI sequence with 5 unweighted measurements and 20 isotropically distributed encoding directions at two non-zero *b-*values of 0.8 and 1.8 ms/μm^2^. Remaining scan parameters were: delta/Delta = 5/11 ms, TE = 21.2 ms and TR = 2000 ms, field-of-view 28 × 28 mm, matrix size 96 × 96, slice thickness 0.6 mm, number of slices 15, number of averages 1). The DKI sequence lasted 36 minutes and total scan time was <1.5 hours. In total, DKI data sets were collected from *n =* 23 animals (11 males and 13 females) in both the awake and anesthetized state.

### 2.6 Postprocessing and DKI data analysis

Prior to analysis all dMRI data were inspected visually for quality. No data sets were discarded. Next, dMRI data were preprocessed in MATLAB (MathWorks Inc, v 2018b). Postprocessing included denoising using NORDIC denoising^29^ and correction for Gibbs ringing^30^. After postprocessing, DKI data analysis was performed in MATLAB using in-house scripts as previously described^24^. From the DKI fits we obtained the diffusion and kurtosis tensors from which MD, FA, and MK were calculated. For further analysis the brains were outlined manually and threshold values for MD and FA were used to segment the brains into GM, WM thus also excluding the ventricles from our analysis. This was done for each data set individually. These masks delineating whole brain, WM, and GM were then used for analysis of MD and MK differences between the awake and anesthetized state.

## 3. Results

### 3.1 Isoflurane anesthesia reduces mouse brain MD and MK in both GM and WM

Visual inspection of whole brain maps of MD and MK estimates for each animal showed subtle changes in MD between awake and anesthesia while the change in MK was more pronounced (**Fig. 2**). Overall, we observed a general reduction in both MD (left column) and MK (right column) for the anesthetized state (bottom row) when compared to awake state (top row). The figure shows maps from the same animal and the presented observations were consistent across all animals (*n* = 23).

**Fig. 2.**
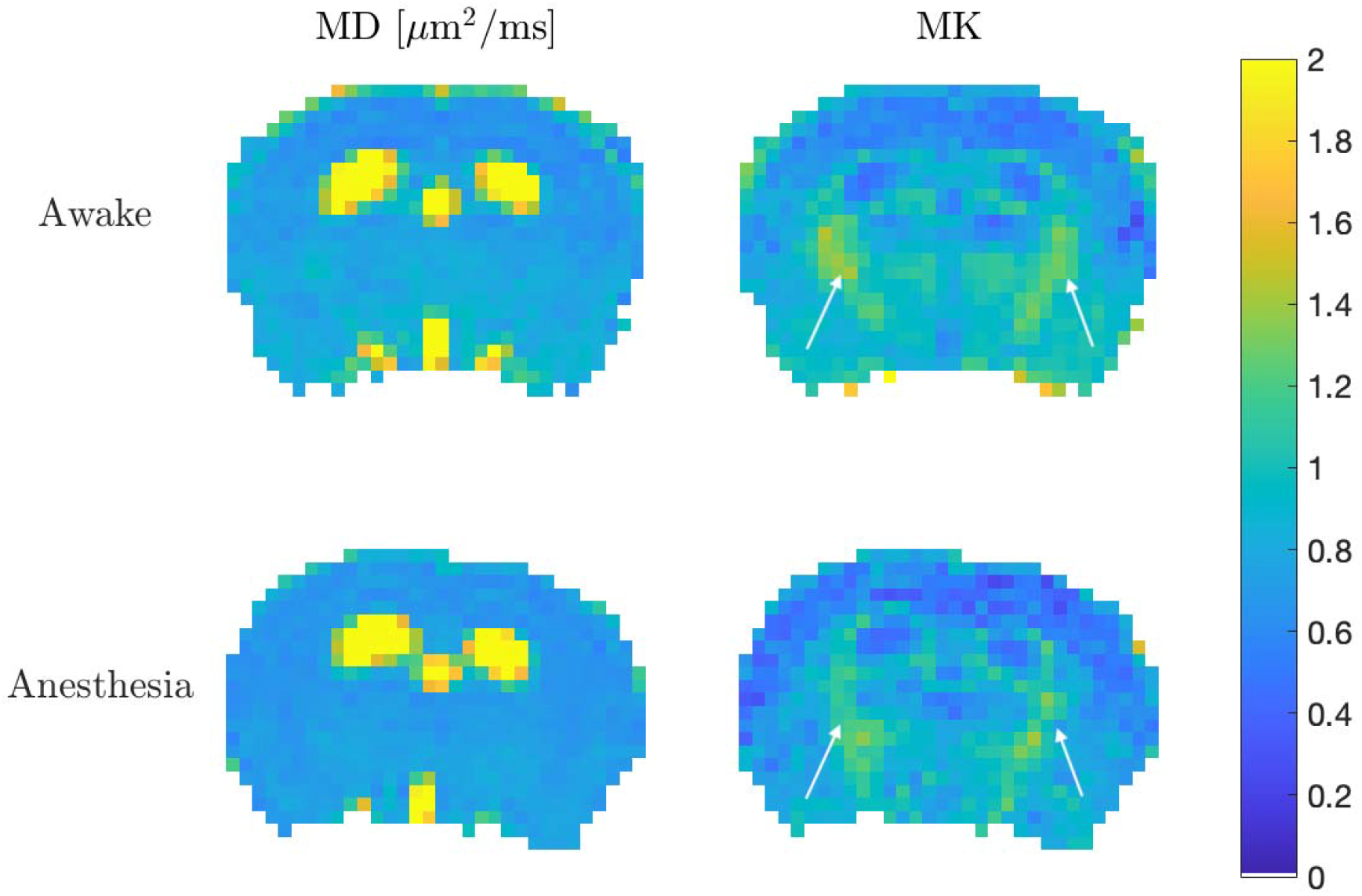
Example maps of MD and MK estimates from a coronal brain section using a representative animal (*n* = 1). A pronounced, overall reduction in MD is seen from awake (left column, top) to anesthetized state (left column, below). White arrows indicate comparable differences in MK estimates of awake and anesthesia states.

Previous findings of anesthesia related ECS volume changes were found in cortical GM tissue, so in our analysis we segment the brains into GM and WM. After segmentation we calculated the average MD and MK in both tissue types and states for each animal. In GM marked changes to both MD and MK is seen between the awake and anesthetized state (**Fig. 3**). We found, that MD in isoflurane-anesthetized mice was decreased compared to awake mice (*P <* 0.05, paired two-sample Student’s t-test) (**Fig. 3, left**), but the effect was somewhat variable among individual mice. In contrast to this the MK change from awake to anesthesia was a significant decrease (*P <* 0.001, paired two-sample Student’s t-test) with uniform response among all mice (**Fig. 3, right**). Collectively, this is indicative that isoflurane anesthesia decreases water diffusivity in the mouse brain and reduces microstructural GM tissue complexity.

**Fig. 3.**
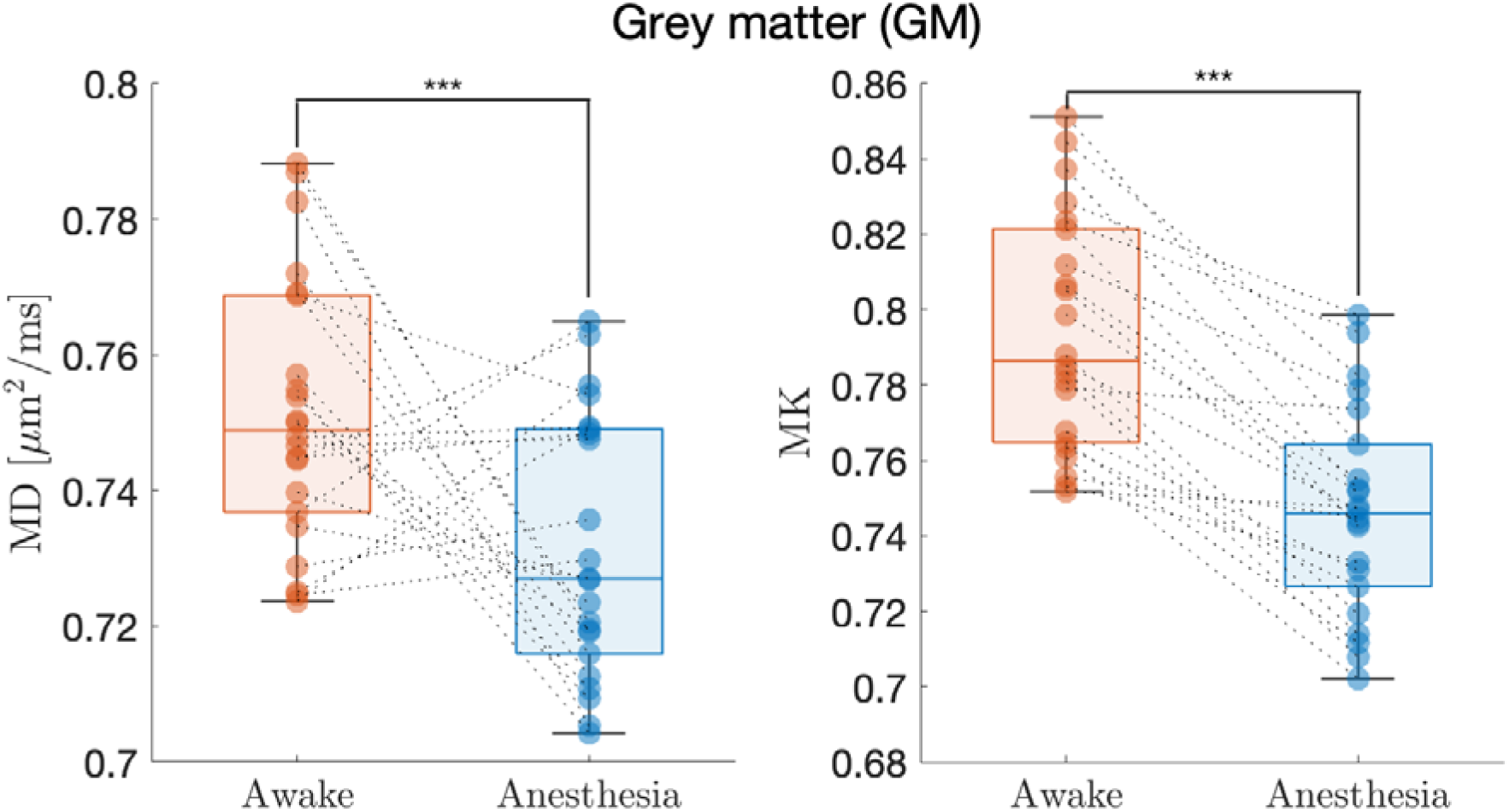
Boxplot with scatter points of each animal’s average GM MD (left) and MK (right) in the awake (orange) and anesthetized (blue) state (*n* = 23). MD is given in units of _μ_m^2^·ms^-1^, MK is unitless. *** Indicates significant difference at the level: *P <* 0.001 (paired two-sample Student’s t-test).

Performing the same analysis for MD and MK in WM we see a similar effect with isoflurane anesthesia significantly decreasing both MD and MK. Also here, a uniform response is seen in MK whereas in MD a few animals exhibit increased MD in the anesthetized state compared to the awake state in opposition to the majority of the cohort.

### 3.1 Simulation results

Our first simulation is based on a generic two-compartment system to illustrate how a shift in the volume fraction of the fast-diffusing compartment (ICS) is reflected in MD and MK. The result is shown in **Fig. 5**.

In the figure it is seen that for such a system a reduction in ICS volume fraction (i.e. an increase in ECS volume) leads to a decrease in MD and an increase in MK for all relevant ICS volume fraction values. While our simulation uses compartmental diffusivities that may not match the exact diffusion properties in mouse brain, we note that the overall behavior observed in **Fig. 5**. does not depend on the distinct values attributed to the two compartments as long as *D*_1_>*D*_2_. Clearly, the behavior observed in **Fig. 5**. shows that an anesthesia related decrease in both MD and MK as seen in our experiments is not produced by ECS expansion in a two-compartment system. To further explore this, we extended our simulations to allow for two independent ICS volume fractions. **Fig. 6**. shows maps of MD and MK from simulations varying the volume fractions of these two compartments. Overall, the addition of a second compartment changes the dynamic drastically compared to the simple response seen in a two-compartment system (**Fig. 5**). Instead, we see here that for a three-compartment model it is possible for a reduction in total ICS volume fraction (*f*_1_+*f*_2_) to produce a decrease in both MD and MK. An example of such a transition is indicated by the arrows in **Fig. 6**. The path produces a decrease in MD from 0.8 µm^2^/ms to 0.7 µm^2^/ms and a reduction in MK from 0.8 to 0.7. These values are seen to be in agreement with the measured values shown in **Fig. 3**. In our simulations, these reductions are produced by a reduction in total ICS volume fraction from approximately 0.8 to 0.6. The basis for these results and their potential physiological implications are covered in the discussion.

## 4 Discussion

Previous dMRI studies were unable to detect significant brain ADC changes between the awake state and isoflurane anesthesia in mouse and between the awake state and natural sleep in humans^15,17^. In this study we set out to investigate if metrics from the microstructurally sensitive DKI method show change between the anesthetized and awake state in brain.

To ensure robust and reliable imaging data from awake mice we habituated our mice to awake MRI as previously described^31^. Following this MRI habituation, we performed DKI scans of mice in the awake state and later in the anesthetized state thereby enabling us to use each mouse as its own control. On the single animal level, no significant changes were observed for either MD or MK in whole brain, WM or GM. On the group level, however, differences in MD and MK were found in both GM and WM. In GM, isoflurane anesthesia significantly decreased mean MD and MK values when compared to the awake state (**Fig.3**). Previous MRI studies have not detected anesthesia/sleep related changes to brain tissue diffusivity^15,17^. We believe that this is caused by previous studies relying on the ADC metric which is estimated from a single encoding direction whereas our MD estimate stems from multiple encoding directions which improves estimation. Furthermore, our study uses the DKI framework which is known to improve MD estimation over DTI-based estimation^25^. We note that even with this improvement in diffusivity estimation our data does show variation in the change in MD with few animals exhibiting instead an increased MD in the anesthetized state. Taken together, these effects may explain why previous studies have not seen an effect. Contrary to MD, the GM MK response was completely uniform with all animals having decreased MK in the anesthetized state compared to the awake state. The same response to anesthesia was seen in WM (**Fig. 4**). Collectively, these findings are indicative of an anesthesia induced microstructural modulation in agreement with the findings by Xie et al.^3^ However, while MK is a sensitive marker for tissue microstructure it is not readily interpretable in terms of tissue compartment sizes. However, it is understood^27^ that for tissue composed of several compartments with different diffusion rates the observed diffusion kurtosis is related to the variance over compartmental diffusivities including a weighting by compartmental volume fractions (Eqs. (4)-(5)). For unchanged compartmental diffusivities a change in MK can therefore be caused by volume fraction shifts alone e.g. an ECS volume increase as reported by Xie et al.^3^ Such an increase must come at the expense of the ICS volume change meaning that cells shrink during sleep and anesthesia. A central question surrounding this mechanism is therefore if all cells shrink or if only a subset of cells modulate their volume during sleep/anesthesia. It also remains to be understood if all cell types shrink to the same degree and if the entire cell shrinks or if only certain cell components modulate their shape/volume as part of this response. It has previously been shown that axon-spine interface on synapses in motor and sensory cortices decreases (roughly 18%) during sleep, however this might be different for other parts of the brain^33^. While sensitive, our DKI methods cannot resolve such details. Nevertheless, we do detect significant changes in WM and GM MD and MK indicative of microstructural modulation during anesthesia. This effect, however, seems to affect all WM and GM equally. In particular, we did not find any GM regions not displaying a reduced MK in the anesthetized state.

**Fig. 4.**
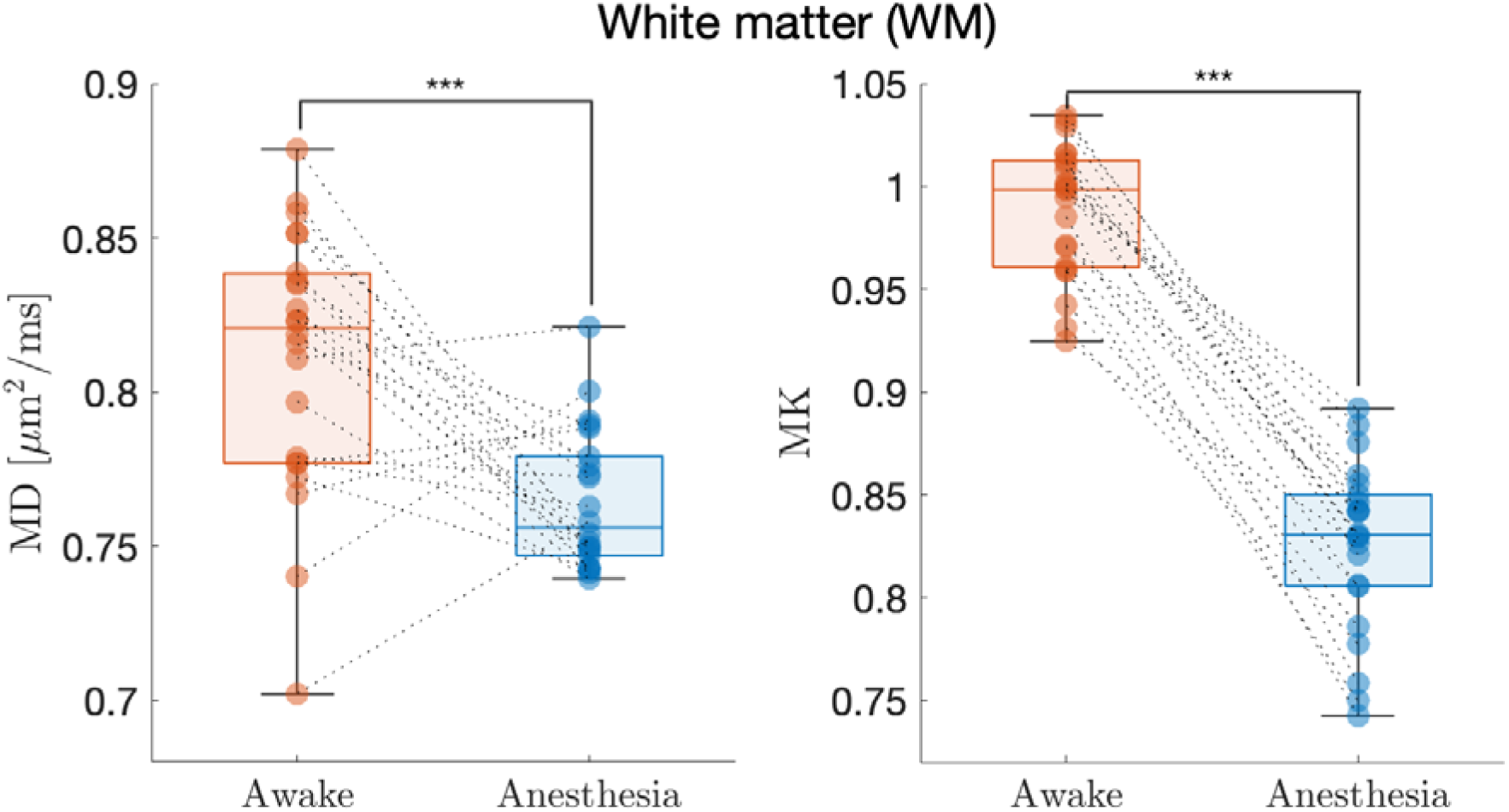
Boxplot with scatter points of each animal’s average WM MD (left) and MK (right) in the awake (orange) and anesthetized (blue) state (*n* = 23). MD is given in units of _μ_m^2^·ms^-1^, MK is unitless. *** Indicates significant difference at the level: *P <* 0.001 (paired two-sample Student’s t-test).

**Fig. 5.**
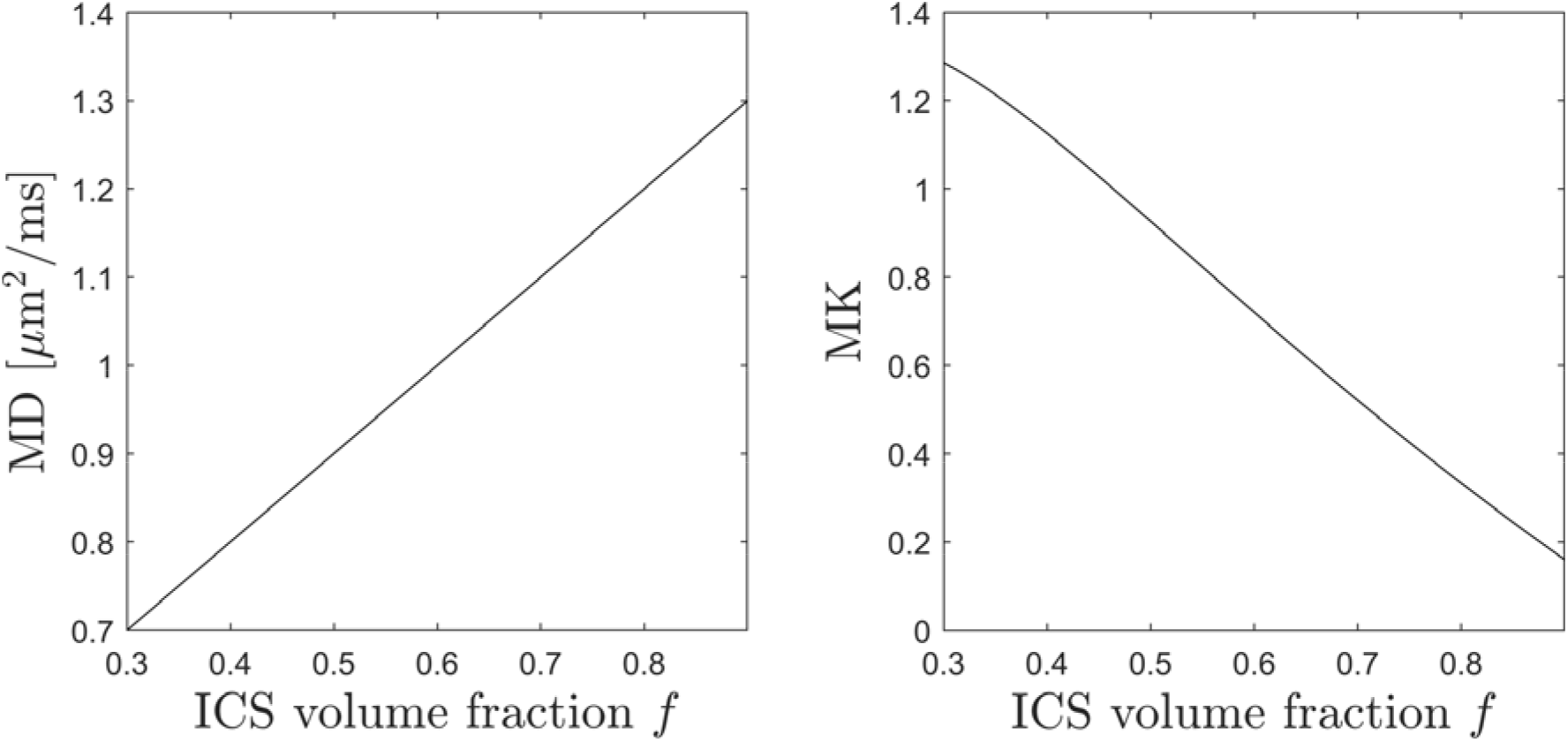
Simulation results from a two-compartment CSF/GM system with *D*_1_ = 1.5 µm^2^/ms (ICS) and *D*_2_ = 0.5 µm^2^/ms (ECS). In both plots the x-axis shows the ICS volume fraction, *f*. The left plot shows the MD produced by such a system as a function of *f*. Similarly, the plot on the right shows the MK from this system.

**Fig. 6.**
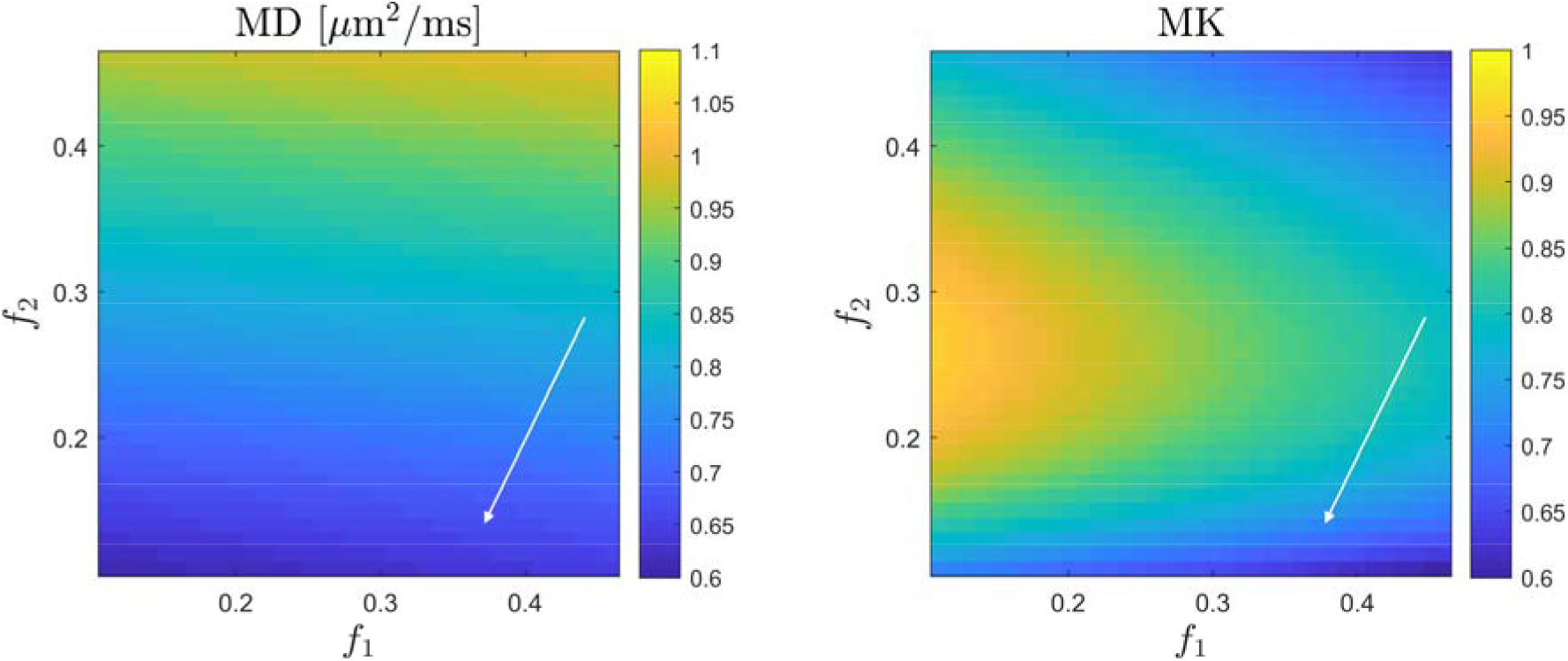
Simulation results from a three-compartment system with two ICS compartments to allow for a differentiated ICS volume response to e.g. sleep or anesthesia. This system has *D*_1_ = 1.5 µm^2^/ms (ICS1), *D*_2_ = 0.6 µm^2^/ms (ICS2), and *D*_3_ = 0.5 µm^2^/ms (ECS). The maps show the MD produced by such a system as a function of the volume fractions *f*_1_ and *f*_2_. The map on the right shows the corresponding MK behavior. The arrows indicate a path where a decrease in total ICS volume *f*_1_+*f*_2_ (i.e. an ECS increase) for such a system can produce a decrease in MD and MK similar to our data.

In our simulations we find that the observed GM MD and MK modulations cannot be achieved with only two compartments. When allowing for three compartments our simulations show that MD and MK can decrease with an ICS volume fraction decrease (corresponding to cell shrinkage or ECS increase). In our simulations, we assign compartmental diffusivity values in agreement with those from previous studies^27,28^ and the relation between ICS/ECS diffusivity (ICS faster than ECS) agrees with microscopy work exploring compartmental diffusivities in fixed and perfused brain samples^34,35^. Our simulations are nevertheless exploratory and serve only to elucidate if a cellular volume fraction change in sleep/anesthesia can be consistent with our DKI observations. The volume fraction changes discussed in relation to the arrows in Fig. 6 are therefore illustrative. Nevertheless, our simulation results may not be entirely unreasonable as recent reports have estimated the ECS volume expansion during sleep and anesthesia to be as much as 60%^3^. Importantly, our simulations indicate that a differentiated cell response seems to be required to explain our finding of decreased MD and MK in the anesthetized state. This is important because known physiology is difficult to reconcile with the ECS expansion observed during sleep and anesthesia^3,12^ if this expansion is produced by shrinkage of all cells. The difficulties with this picture relate to the functions of the various brain cell types. Neurons depend critically on correct ion concentrations for their function and therefore such volume change would have to be a closely controlled process for neurons to retain their function during such sleep/anesthesia-induced shrinkage^3^. Similarly, glia cells in general and pericytes, in particular, have been assigned new, important roles in recent years. In the brain, pericytes are essential for maintaining the structural integrity of the blood-brain barrier (BBB) and for allowing the entry of immune cells into the brain^36,37^. Furthermore, due to their contractility pericytes are believed to have an important role in controlling the brain’s blood flow^38–40^. All of these functions seem difficult to reconcile with a sizeable cell volume reduction during sleep or anesthesia. Consequently, if the ECS volume increases during sleep and anesthesia while global MD and MK is decreased, then it suggests that this must occur based on either a differentiated shrinkage among cell types or an orchestrated cell shrinkage so that not all brain cells shrink at the same time.

The insight offered by our data and simulations is valuable as it is in agreement with previous optics-based observations of anesthesia related microstructure modulations. This implies, that DKI can be used to non-invasively study this phenomenon further. Importantly, such studies can be undertaken in both animals and humans and with whole brain coverage. More advanced analysis and simulation work may be able to more fully explain the tissue dynamics producing our observations. Such work, however, is beyond the scope of this paper where we are focused on demonstrating the effect using well-established methods.

### 4.1 Translational value of understanding ECS dynamics

Several contemporary techniques are well-suited for the exploration of the diffusion properties in ECS and the dynamics during sleep or anesthesia^55^. While animal studies using TMA^+^ quantification provide direct assessment of ECS volume and complexity^3^, it is nonetheless measurements stemming from a small cortical area. Here, DKI is particularly well-suited for whole brain study and is easily applied to humans as well^17,21,56,57^. This avenue of research is very important for our basic understanding of the physiological role of sleep and its connection to brain waste clearance. The ECS remain one of most unexplored areas of neuroscience^55^, but it has nonetheless important features in brain homeostasis and may also provide a new mode of drug delivery to CNS. This could potentially be used with emerging techniques for opening of the BBB, such as focused ultrasound^58,59^ and noninvasive, focused shockwave (FSW) pulses^60^. However, drug delivery to the brain across BBB via intrathecal pathways is far from well understood and it remains for future studies to fully characterize particles of different sizes, their properties and how these impact the effectiveness of delivery in combination with ECS dynamics. In-depth understanding of anesthetic or sleep induced ECS dynamics is essential for the developmental success of these therapeutic interventions.

### 4.2 Effect of anesthesia and implications for preclinical dMRI

Our study compares data obtained in anesthetized mice to the awake state in the same animals. Admittedly, MRI studies comparing dMRI data from the awake state to data obtained during natural sleep would have been preferable. However, due to the loudness of MRI scans in general and dMRI in particular this was not possible to achieve. In Xie et al.^3^ an ECS volume increase was observed in natural sleep and the anesthetized state with very similar effect between both states. However, in their study, a ketamine-xylazine mixture was used for anesthesia maintenance, which may not produce effects directly comparable to the gas isoflurane anesthesia used in the present study. Although the molecular mechanisms of isoflurane and ketamine-xylazine are well known, the details of how neurophysiology and cardiovascular function is altered between the different anesthetics is not well-known. Comparisons should therefore always be made with caution. For example, isoflurane anesthesia has been shown to induce vasodilation which leads to increases in coronary and cerebral blood flow (CBF)^41,42^, an effect that is not observed in rodents during ketamine-xylazine anesthesia^43^. Although CBF measurements were not conducted in our study, it is fair to assume that CBF are altered between the awake and anesthesia state in our study. However, we do not consider this effect something that influences our dMRI data in any significant way and certainly not to the extent, that it can be said to be the cause of the WM and GM diffusion changes observed in our study. Firstly, the effect we observe is most robust in MK which is estimated using high b-values (0.8-1.8 ms/μm^2^) unaffected by intravascular incoherent motion (IVIM) effects^44,45^ and in tissue with low vascular volume fraction (GM blood volume is estimated to be approx. 2-5%^46^). Secondly, our analysis uses the magnitude images whereas any flow effect would only manifest in the phase images.

Besides the obvious CBF alterations of isoflurane there are also other important aspects when comparing different anesthesia regimes (i.e. isoflurane and ketamine-xylazine) and natural sleep in the context of our findings. In general, anesthesia serves as an excellent tool for inducing unconsciousness and may share many neurobiological processes with natural sleep^47–49^. However, anesthesia is not equivalent to natural sleep^50^ and comparisons of these states are therefore inevitably flawed. Even though, different anesthetic agents can resemble the physiological state of sleep more closely than others^51^. For example, EEG measurements in and mice and rats have shown that ketamine-xylazine produces synchronized, slow-oscillatory activity similar to what can be observed during slow-wave sleep^3,51^. In contrast, in rats and humans isoflurane anesthesia produces patterns of burst-suppression mimicking a state of coma suggesting that ketamine-xylazine produces a physiological state closer to natural sleep when compared to isoflurane^51–53^.

Interestingly, rodents under ketamine-xylazine anesthesia have been shown to have greatly increased CSF transportation compared to isoflurane anesthetized animals^54^. It is not clear how the CSF transport rate affects GM/WM microstructure but these reports do suggest that characterization of DKI parameters in ketamine-xylazine anesthesia would be of interest to the field. Ongoing work in our group investigates the effect of different types of anesthesia on DKI derived parameters as well as temporal aspects of the microstructural modulation observed in our data using the fast-DKI method^61^. Since the simulations from this study indicate that cell types may respond in a differentiated manner during the transition to anesthesia, we are aiming to explore this phenomenon with a combination of optical methods, such as two-photon microscopy, and MRI to provide a better foundation for improved interpretation^62,63^.

While the choice of anesthetic in this study must be discussed when comparing our findings to other studies (in particular Xie et al.) it must be noted that our findings have relevance outside of the glymphatics research field. Specifically, we note that isoflurane is the most widely used anesthetic for *in vivo* rodent MRI. The observation that brain microstructure in anesthetized animals differs from their value in the awake animal (and therefore in the natural state of the brain) is important for the whole field of preclinical MRI. These findings further underscore the scientific value of the transition to awake animal MRI.

## Conclusion

We report a difference in both MD and MK between the awake and the anesthetized mouse brain. The effect is observed in both GM and WM. Our findings are consistent with the reports of brain ECS volume increase in sleep and anesthesia. We use simple simulations to interpret our findings and find that a differentiated cellular response is needed for the DKI changes we observe potentially reconciling an ECS volume increase during sleep and anesthesia with other aspects of brain physiology. Our study shows that DKI has potential as a tool for the study of brain microstructure alterations during anesthesia and sleep in both animals and humans. Furthermore, our study demonstrates an overlooked effect of anesthesia on brain microstructure which is relevant to preclinical MRI as a whole.

